# RM1mAb avoids Intracellular degradation to Synergistically inhibit NLRP3 Inflammasome Activation in Familial Cold Autoinflammatory Syndrome

**DOI:** 10.64898/2025.12.09.693294

**Authors:** Angela Lackner, Sofia I. Picucci, Valerie Henriquez, Karen Wang, Reginald McNulty

## Abstract

The NLRP3 inflammasome enables release of mitochondrial DNA to circulation. Circulating oxidized mitochondrial DNA generated in response to NLRP3 inflammasome activation functions as an alarmin that contributes to the maintenance of systemic inflammation. The discovery that NLRP3 could cleave oxidized mtDNA led to repurposed chemical inhibitors that dually target NLRP3 and DNA glycosylase OGG1, resulting in pro-survival type-1 interferon. Using molecular dynamics and immunology we show that RM1mAb, a full-length monoclonal antibody targeting the NLRP3 pyrin domain, can prevent NLRP3 from interacting with mitochondrial DNA. We further illustrate RM1mAb can exploit FCγRs for cell entry and avoid destruction by the lysosomal pathway to inhibit IL-1β secretion in peripheral mononuclear blood cells (PBMCs) isolated from patients with Familial Cold Autoinflammatory Syndrome harboring NLRP3 L353P gain of function mutation. We show RM1mAb and repurposed inhibitor TH5487 synergistically inhibit inflammasome activation. These findings illustrate the promise of exploiting FCγRs as a means of IgG entry to target cytosolic proteins and improve human health.

**One-Sentence Summary:** RM1mAb and TH5487 synergistically inhibit inflammasome activation in human FCAS PBMCs.

## Introduction

Circulating mitochondrial DNA is linked to inflammaging and is a sign of mortality (*1, 2*). NLRP3 inflammasome activation is linked to extracellular release of oxidized mitochondrial DNA to the blood stream which fuels inflammation in a plethora of diseases including sepsis (*3*), myelodysplastic syndromes (MDS)(*4*), and systemic lupus erythematosus(*5*). NLRP3, which binds and cleaves oxidized mitochondrial DNA(*6–9*), can have these interactions and activation simultaneously inhibited with repurposed drugs that also target human glycosylase OGG1 which has a similar protein fold to NLRP3 pyrin domain(*7*).

Currently, there are no FDA-approved direct NLRP3 inhibitors(*10*). The most well-known direct NLRP3 inhibitor, MCC950, showed liver toxicity in a phase II clinical trial and was removed from development and ineffective in patients with hyper-active NLRP3 containing the L353P mutation(*11*). Additional efforts have focused on systemically inhibiting interleukin-1β (IL-1β) downstream of NLRP3 activation. Canakinumab, a monoclonal antibody against IL-1β that prevents receptor interaction(*12*) is FDA-approved for treating CAPS. Anakinra is an IL1-receptor antagonist that prevents IL-1α/β interaction(*13*) is also FDA-approved for CAPS. These drugs are not direct NLRP3 inhibitors but inhibit downstream cytokine production from several inflammasomes including NLRC4, AIM2, pyrin, NLRP1(*14*) and inflammasome-independent pathways such as Caspase-8-dependent IL1β processing(*15*).

Herein, we report development of a full-length monoclonal antibody that enters the cell via FCγR1 that avoids lysosomal degradation to target NLRP3 pyrin domain to inhibit inflammasome activation in peripheral mononuclear cells from patients with Familial Cold Autoinflammatory Syndrome (FCAS) L353P(*16*), a subcategory of Cryopyrin-Associated Periodic Syndrome (CAPS). Additionally, this antibody inhibits NLRP3 from interacting with mtDNA. These results have implications ameliorating increases in circulating mitochondrial enabled by NLRP3 inflammasome activation in systemic inflammatory diseases.

## Results

### *In-Silico* generation and validation of a humanized monoclonal antibody targeting the NLRP3 pyrin domain

Small molecule drugs like MCC950 have been designed to target NLRP3 directly via the NACHT domain(*17*). Unfortunately, this drug has severe off-target effects and memetics based on MCC950 have been made to no avail(*10*). Recent progress by the McNulty lab illustrated repurposed inhibitors binding the pyrin domain could inhibit human glycosylase OGG1 and NLRP3 inflammasome activation(*7*). Nonetheless, we sought to make a specific monoclonal antibody-based inhibitor to target the NLRP3 pyrin domain. Our first step was to design an entire IgG1 *in-silico* involved making the antigen binding domains known as the variable heavy (VH) and variable light (VL) domains, and subsequently grafting those onto a human IgG backbone containing the constant region of the light chain, and the CH1, CH2, and CH3 domains of the heavy chain, to create a finalized design of two Fab regions and a FC region (**Fig. 1A**). We utilized RFantibody pipeline to design and optimize the variable regions directed to the NLRP3 pyrin domain(*18*). The 3D models of the variable regions were inspected for proper folding using AlphaFold3(*19*). We selected 2 out of 30 VH/VL pairs (denoted herein as RM1mAb and RM2mAb) predicted to bind directly to the NLRP3 pyrin domain (**Fig. 1B, C**). We then grafted these variable regions onto the constant regions of the human antibody from PDB 1HZH(*20*) using PyROSETTA(*21*). To further verify proper folding, we simulated the antibody structure using AlphaFold3 (**Fig. 1D, E**). These constructs were cloned and expressed in Expi293 cells and purified with low contamination using protein A affinity chromatography to approximately 0.5 mg/mL (**Fig. 1F**).

**Figure 1:**
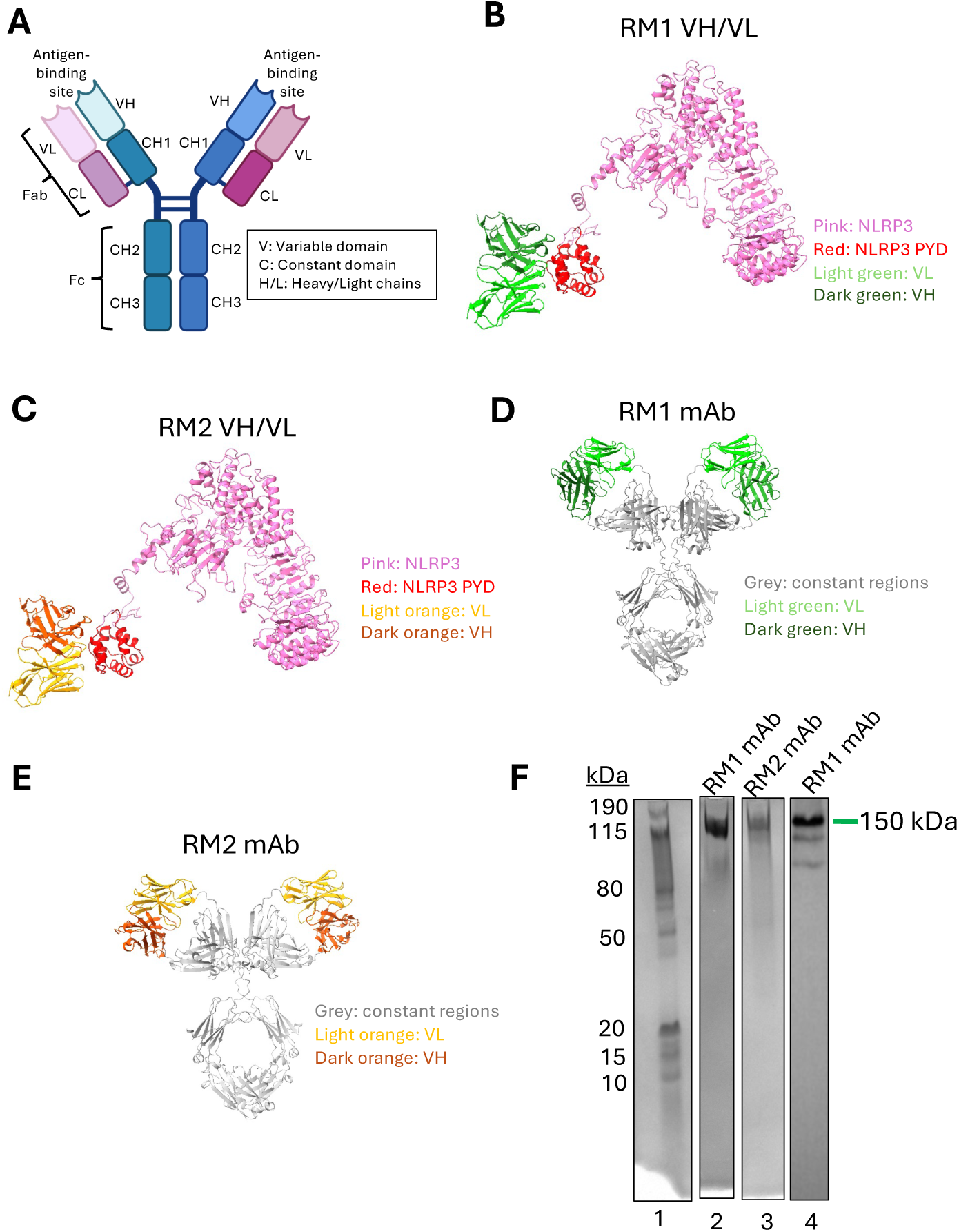
Design and purification of anti-PYD mAbs. A) Key aspects of an antibody to include in our designs. AlphaFold3 predictions of the variable heavy (VH) and light (VL) chains of mAbRM1 (B) and RM2 (C) binding to NLRP3 pyrin domain (PYD). AlphaFold3 predictions for folding of RM1mAb (D) and RM2 (E). F) Homogenous purification of RM1 and RM2. Lane 1: Molecular weight ladder. Lane 2: SDS Page gel of RM1mAb. Lane 3: SDS Page gel of RM2. Lane 4: Anti-human western blot.

To examine the integrity of the NLRP3-PYD-mAb complex, we sought to examine the stability of the interface with nanosecond timescale all-atom conventional MD simulations with NAMD3(*22*). The final system comprised of 25958 atoms and solvated with the TIP3P explicit water molecules with CHARMM36(*23*) force field for 50 ns (**Supplemental Movie 1**). The complex interface maintained average of 1322 contacts within 5 Å and 7 hydrogen bonds throughout the simulation (**Fig. 2A**). Initial inspection of torsion *phi* and *psi* angles contained 98% in favorable regions 99.8% residues in allowed regions (**Fig. S1A, B**). The conformational landscape of NLRP3 revealed multiple discrete metastable states, and RM1mAB showed a similar pattern, sampling high- and low-RMSD conformations that closely tracked movements at the binding interface (**Fig. S1C, D**). A 2D conformational density map further demonstrated that NLRP3 repeatedly revisited specific high-probability structural substates rather than undergoing continuous drift. Together, these results indicate coordinated yet stable flexibility within the complex, likely arising from the inherent mobility of the flexible linker connecting the pyrin and NACHT domains (**Fig. S1E**).

**Figure 2:**
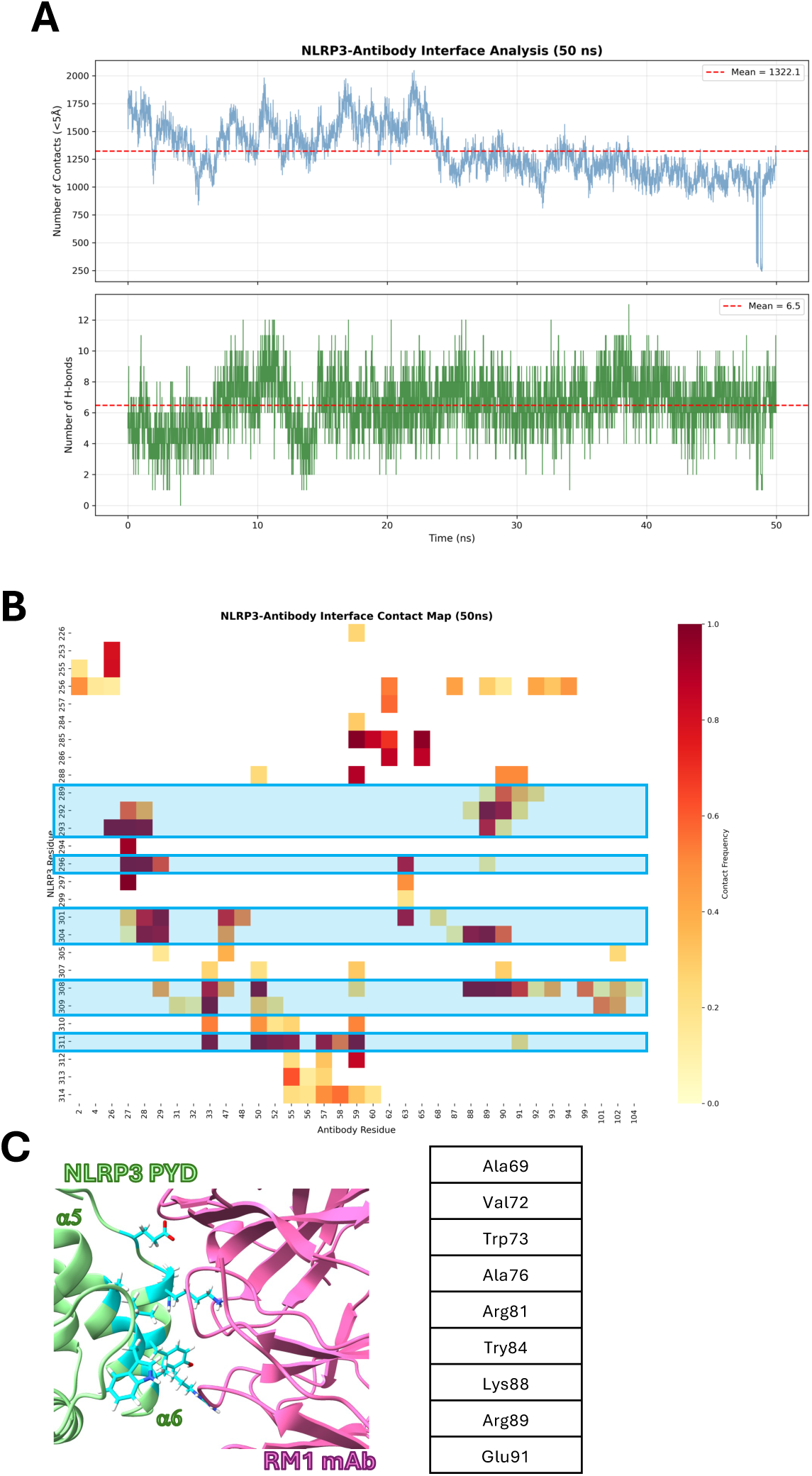
Molecular dynamics reveals significant contacts and hydrogen bonds are formed between NLRP3 and RM1mAb. A) Analysis of <5A contacts (top) and hydrogen bonds (bottom) that occur between NLRP3 and RM1mAb between 1 and 50 ns. B) The interface contact map of per-residue contacts between NLRP3 and RM1mAb between 1 and 50 ns. Blue boxes correspond to top contacts shown in (C) C) Top contact residues (cyan) from NLRP3 (green) to RM1mAb (pink) identified in (B) shown both visually (left) and in a table (right).

The NLRP3-RM1mAb per-residue interface was represented using a contact heat map (**Fig. 2B**). We identified 27 residues in NLRP3 and 35 residues in RM1mAb that had a role in this interface (**Fig. 2B**) (**Table S1**). Also, several NLRP3 residues at the interface with RM1mAb that illustrated high-contact frequency included A69, V72, W73, A76, R81, Y84, K88, R89, and E91. These amino acids are all located on the fifth and sixth helices of the NLRP3 pyrin domain (**Fig. 2C**). In all, many NLRP3 residues between 33-94 had some sort of interaction with RM1mAb. The antibody did not interact at all with the NACHT or LRR domain. Altogether, these data illustrate our *in-silico* interactions should hold up with *in vitro*. We sought to move forward with RM1mAb to examine efficacy.

### Anti-pyrin mAb directly binds NLRP3 and prevents NLRP3 from binding mitochondrial DNA

NLRP3 bound to oxidized DNA is not recognized by monoclonal pyrin-targeting antibodies against the NLRP3 pyrin domain. Conversely, NLRP3 bound to a pyrin targeting mAb, does not bind oxidized DNA(*6*). This suggest NLRP3 pyrin domain interacts with a mAb in a similar fashion to oxidized mitochondrial DNA. Therefore, we examined if there were similar interactions *in silico* between NLRP3 pyrin and oxidized DNA and our newly designed RM1Ab using AlphaFold. We superposed the pyrin domains with models of each and found the interaction surface to be essentially the same (**Fig. 3A, B**). Pyrin superposition resulted in clashes between the oxDNA and VH/VL regions of RM1mAb (**Fig. 3C**). Amino acids of NLRP3 pyrin interacting within 5 Å of both RM1mAb and ox-mtDNA included residues W73, A67, and R81. This suggests binding of one might inhibit binding of the other, as they are competing for the same site.

**Figure 3:**
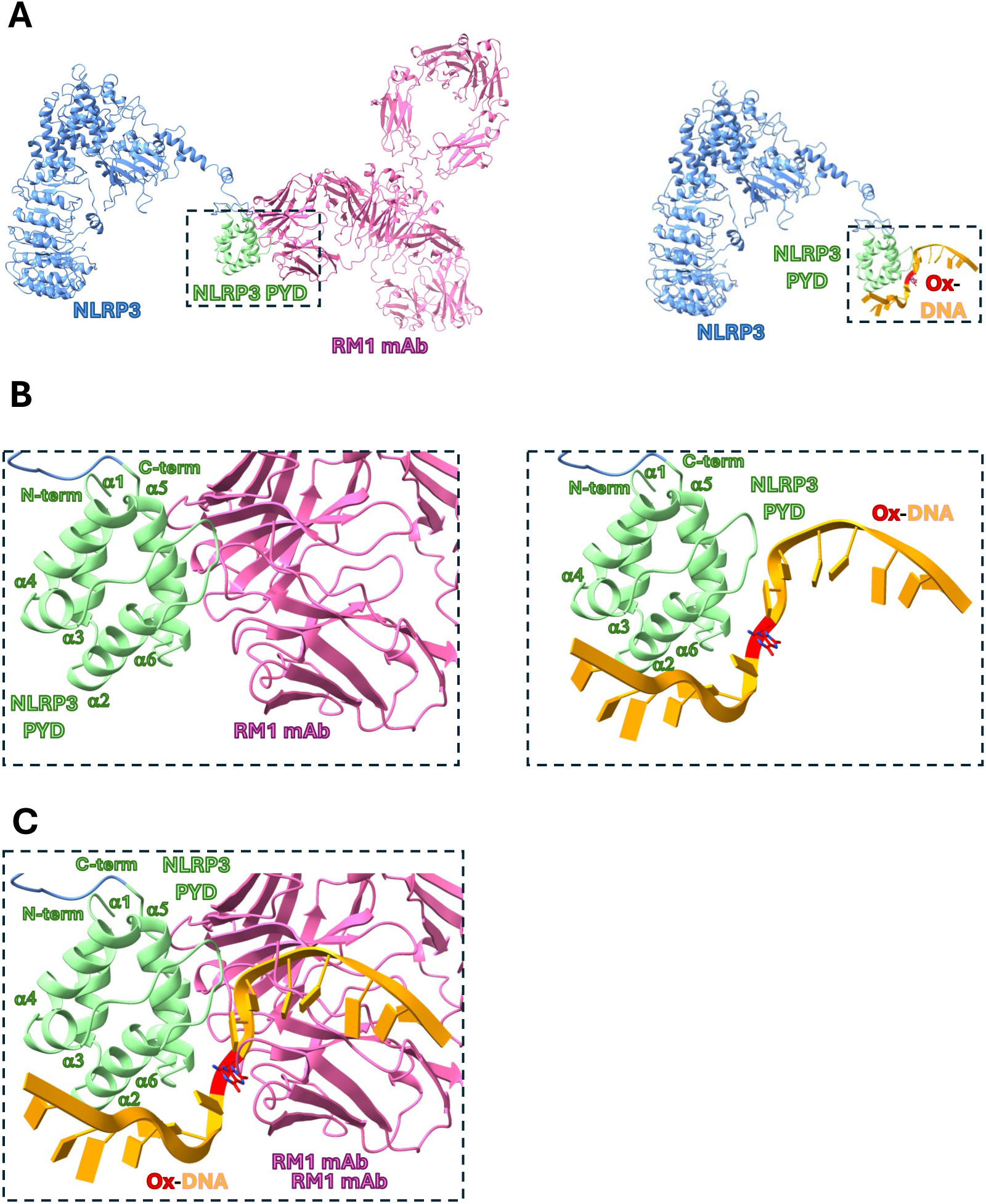
Oxidized DNA and RM1 mAb bind in the same location on the NLPR3 PYD. A) AlphaFold simulation of the association between the NLRP3 (blue) PYD (green) and RM1mAb (pink) (left) and NLRP3 and oxidized (red) DNA (orange) (right) B) Zoomed in box from (A) highlighting the binding interface between NLRP3 and RM1mAb (left) and NLRP3 and oxDNA (right). C) The binding interface to NLRP3, RM1mAb, and oxDNA are all shown together to highlight the clashing interfaces.

Electromobility shift assay is typically used to analyze protein-DNA interactions(*24*). However, a supershift can be obtained with the protein-DNA complex is also bound to an antibody(*25*). As proof of concept that RM1mAb could bind NLRP3, we performed EMSA-like assay under native non-denaturing conditions. RM1mAb migrated higher in the gel in the presence of NLRP3 compared to RM1mAb alone (**Fig. 4A**). We confirmed this using an ELISA assay showing NLRP3 binds directly RM1mAb when the antibody is immobilized on immunoblot plates (**Fig. 4B**).

**Figure 4:**
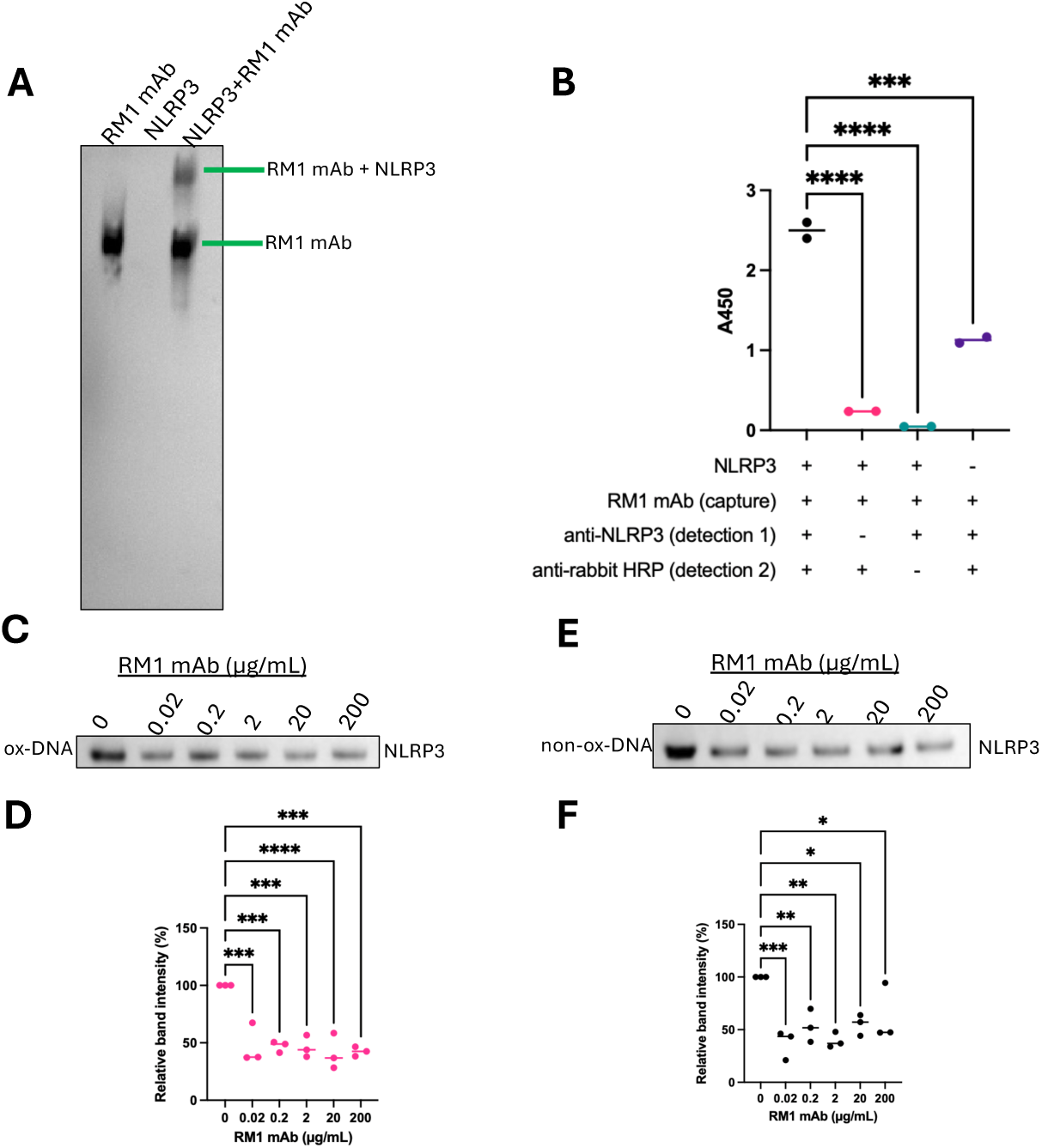
RM1 mAb binds directly to NLRP3 and specifically inhibits interaction with DNA. A) EMSA shows a size shift of RM1mAb by western blot when incubated with NLRP3. B) Microtiter plates were coated with RM1mAb and incubated with or without NLRP3. They were then incubated with or without and anti-NLRP3 Rb primary detection antibody, then with an anti-Rb HRP linked secondary detection antibody. A chemiluminescence reaction was performed and the A450 value of signal intensity was read by plate reader. N=2, Analyzed by one way ANOVA where ***P<0.001 and ****P<0.0001 C) NLRP3 was preincubated with various concentrations of RM1mAb before being incubated with immobilized oxidized DNA. The amount of NLRP3 that bound to the immobilized DNA was visualized by western blot. D) Quantification of the band intensity from (C) showing RM1mAb inhibited NLRP3 binding to oxidized DNA with as little as 0.02 ug/mL. E) NLRP3 was preincubated with various concentrations of RM1mAb before being incubated with immobilized non-oxidized DNA. The amount of NLRP3 that bound to the immobilized DNA was visualized by western blot. F) Quantification of the band intensity from (E) showing RM1mAb inhibited NLRP3 binding to non-oxidized DNA with as little as 0.02 ug/mL. (C-F) N=3, error bars signify mean +/- SEM. Analyzed by one way ANOVA where *P<0.05, **P<0.01, ***P<0.001, ****P<0.0001. G)) NLRP3 was preincubated with various concentrations of RM1mAb, or a non-specific Rabbit IgG, before being incubated with immobilized oxidized DNA. The amount of NLRP3 that bound to the immobilized DNA was visualized by western blot.

Next, we performed a competition assay to see if NLRP3 pre-incubated with RM1mAb prevented NLRP3 from interacting with mitochondrial DNA. RM1mAb prevented NLRP3 from interacting with both oxidized and non-oxidized mitochondrial DNA-coated Dynabeads. The amount of NLRP3 remaining on the beads in the presence of RM1mAb decreased 62.7% and 56.5%, for oxidized and non-oxidized mtDNA, respectively (**Fig. 4C-D**). Preventing NLRP3 interaction with mitochondrial DNA is significant since the direct interaction is sufficient for inflammasome activation.

### NLRP3 monoclonal antibody inhibits inflammasome activation

We previously showed drugs that inhibit NLRP3 interaction with oxidized mitochondrial DNA also inhibit inflammasome activation(*7*). Since RM1mAb binds NLRP3 and prevents interaction with mitochondrial DNA, we examined if RM1mAb could inhibit inflammasome activation. A major hurdle for an antibody to inhibit cytosolic targets is its ability to enter the cell (**Fig. 5A**). To examine if anti-NLRP3 pyrin IgG could be absorbed by cells thereby utilizing endogenous Fc receptors (FcRs), we fluorescently labeled the mAb lysines with AlexaFlour 640, treated THP1 cells by treatment with or without electroporation, and measured localization of the fluorophore with the cells. We showed that both treatment strategies resulted in cellular absorption of the antibody (**Fig. S2A**). We first tried electroporation of a monoclonal IgG (anti-NLRP3 pyrin IgG) we previously reported could not bind NLRP3 when NLRP3 was bound to oxidized mtDNA (*7*). Electroporation of this antibody followed by LPS/ATP activation inhibited IL-1β secretion in a dose dependent manner (**Fig. 5B**). We found 0.01 μg/ml and 0.1 μg/ml anti-NLRP3-pyrin IgG caused 44.3% and 89.3% decrease in IL-1β secretion, respectively (**Fig. 5C**).

**Figure 5:**
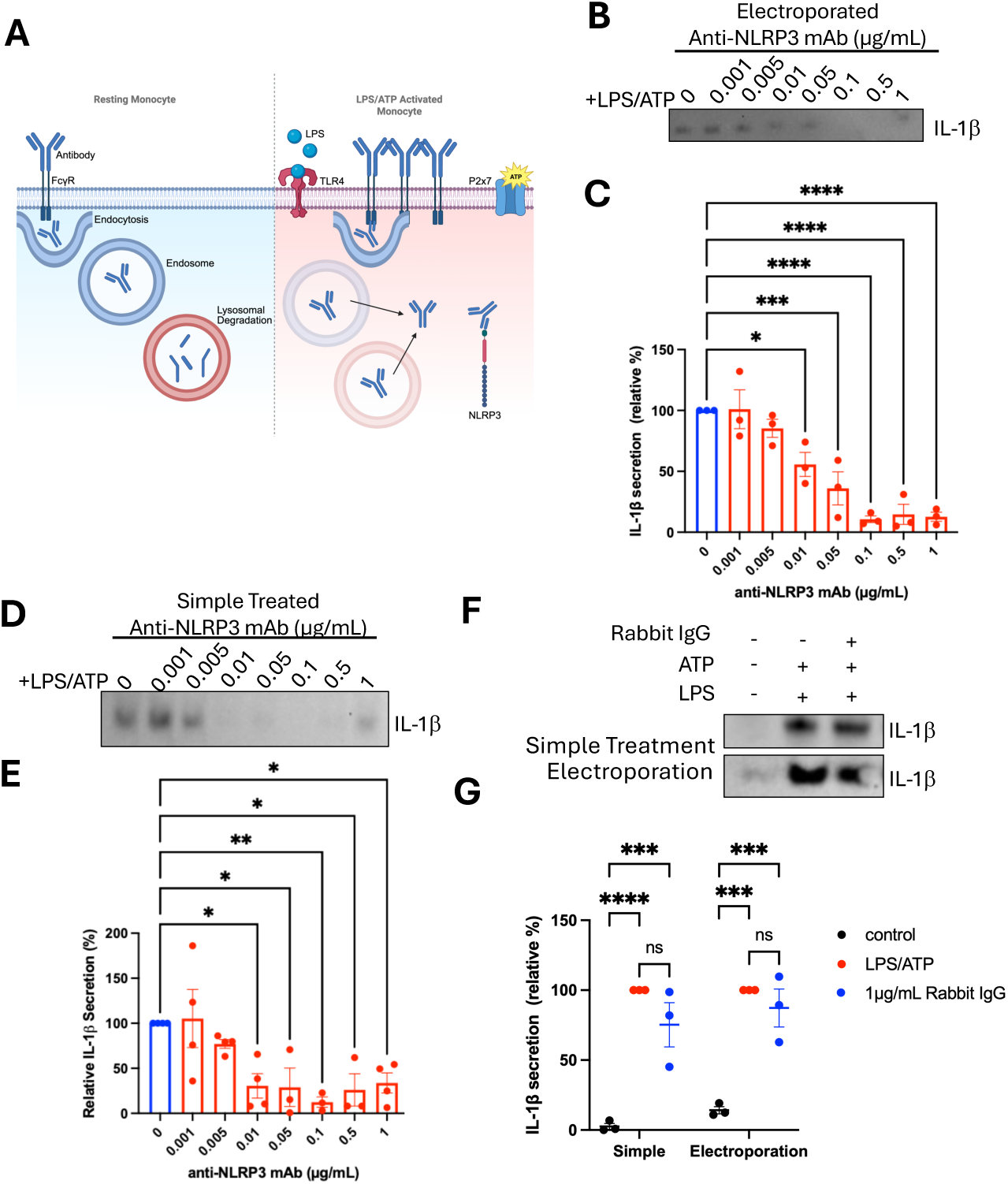
NLRP3-dependent IL-1β release can be inhibited by treating human THP1 cells with anti-NLRP3 antibodies. A) Schematic representation of the internalization of antibodies into cells through Fc receptors in the context of healthy/resting monocytes (left) and LPS/ATP activated monocytes (right). B) Immortalized human THP1 cells were primed with LPS, challenged with 0.001-1 µg/mL Anti-NLRP3 mAb, and activated with ATP. IL-1β secretion was visualized by western blot. C) IL-1β secretion was quantified by evaluating western blot band intensity, showing inhibition with 0.01 ug/mL anti-NLRP3 mAb. D) Immortalized human THP1 cells were electroporated with anti-NLRP3 mAb at 0.001-1 µg/mL, then primed with LPS and activated with ATP. IL-1β secretion was visualized by western blot. E) IL-1β secretion was quantified by evaluating western blot band intensity, showing inhibition with 0.01 ug/mL anti-NLRP3 mAb. F) Immortalized human THP1 cells were primed either treated by simple treatment or electroporation with 1 µg/mL of a normal Rabbit IgG. The cells were also primed with LPS then activated with ATP. IL-1b secretion was visualized by western blot. G) IL-1β secretion was quantified by evaluating western blot band intensity. For all graphs, N=3, error bars signify mean +/- SEM. Analyzed by one way ANOVA where *P<0.05, **P<0.01, ***P<0.001, ****P<0.0001

Cell stimulation under conditions of inflammasome activation causes an upregulation of FcγR1 (Fc-gamma receptor 1)(*26, 27*). IgG has high affinity for these receptors in monocytes and macrophages (*28*). IgG binding to FcγRs can result in antibody internalization, however the fate of the internalized antibody is either lysosomal destruction(*29*) or recycling back to the cell surface (*30*). Interestingly, cellular conditions caused by agents that activate inflammasome activation cause a decrease in functional aspects of the lysosomal pathway(*31–33*) and upregulation of FcγRs (*26, 27*). Therefore, we examined if a full IgG targeting the NLRP3 pyrin could inhibit inflammasome activation in the absence of electroporation. We found dose-dependent inhibition of NLRP3 in stimulated cells without using electroporation (**Fig. 5D**). A dose of 0.01 μg/ml resulted in 69.4% inhibition. Inflammasome inhibition using this method was 36.2% more effective at 0.01 μg/ml compared to electroporation alone (**Fig. 5E**). Inflammasome activation in the presence of a non-targeting rabbit IgG control had no effect on IL-1β secretion, confirming that effects of RM1mAb are NLRP3-specific (**Fig. 5F, G**).

We detected a 221% and 224% increase in pro-caspase-1 retention with treatments of 0.05 and 1 µg/mL antibody, respectively (**Fig. S2B,C**). Importantly, we also confirmed that neither electroporation alone, in the presence of an NLRP3 targeting or non-targeting antibody, of the simple treatment of an anti-NLRP3 antibody induced significant cell death compared to controls (**Fig. S2D-F**)

### Small molecule antibody cocktail synergistically inhibits inflammasome activation in CAPS

Next, we examined if RM1mAb could also inhibit inflammasome activation in human monocyte THP-1 cells. We found 0.5 μg/ml and 1 μg/ml RM1mAb inhibited inflammasome activation by 66.8% and 81.6%, respectively (**Fig. 6A, B**). We detected a 6.3-fold and 6.9-fold increase in pro-caspase-1 retention with 0.5 and 1 µg/mL RM1mAb, respectively (**Fig. S3 A-B**). Analysis of RM1mAb in peripheral blood mononuclear cells (PBMCs) stimulated with LPS/ATP with 0.005 μg/ml and 1 μg/ml illustrated a decrease of 24.1% and 34.8% respectively (**Fig. 6C, D**). A dose of 0.1-1 µg/mL RM1mAb improved cell viability (**Fig. S3C**). Then we tested RM1mAb on PBMCs isolated from CAPS patients with Familial Cold Autoinflammatory Syndrome (FCAS) containing the hyperactive NLRP3 mutation L353P, stimulated with LPS. RM1mAb with a dose of 0.05 μg/ml resulted in 13.8% inhibition, where treatment with the NACHT-targeting NLRP3 inhibitor MCC950 had no effect (**Fig. 6E**). These treatments did not induce significant cell death compared to controls (**Fig. S3D**). Then we examined if we could obtain synergistic inflammasome activation inhibition using doses of RM1mAb and TH5487(*34*) that did not inhibit inflammasome activation. We found 0.01 μg/ml RM1mAb and 5 μM TH5487, doses that on their own do not inhibit, synergistically reduce inflammasome activation by 24.3% (**Fig. 6E**). TH5487, a repurposed DNA glycosylase 1 (OGG1) inhibitor that also inhibits NLRP3, is over-expressed in mitochondria and involved repair of 8-oxo-dG containing mtDNA(*35*). In theory, solely inhibiting OGG1 would promote NLRP3 inflammasome activation. But because TH5487 also targets cytosolic NLRP3(*7, 10, 34*), inflammasome activation is inhibited. Dually targeting NLRP3 with chemical TH5487 and biological RM1mAb illustrates a promising mechanism to decrease systemic inflammation in inflammasome-sensitive disease contexts.

**Figure 6:**
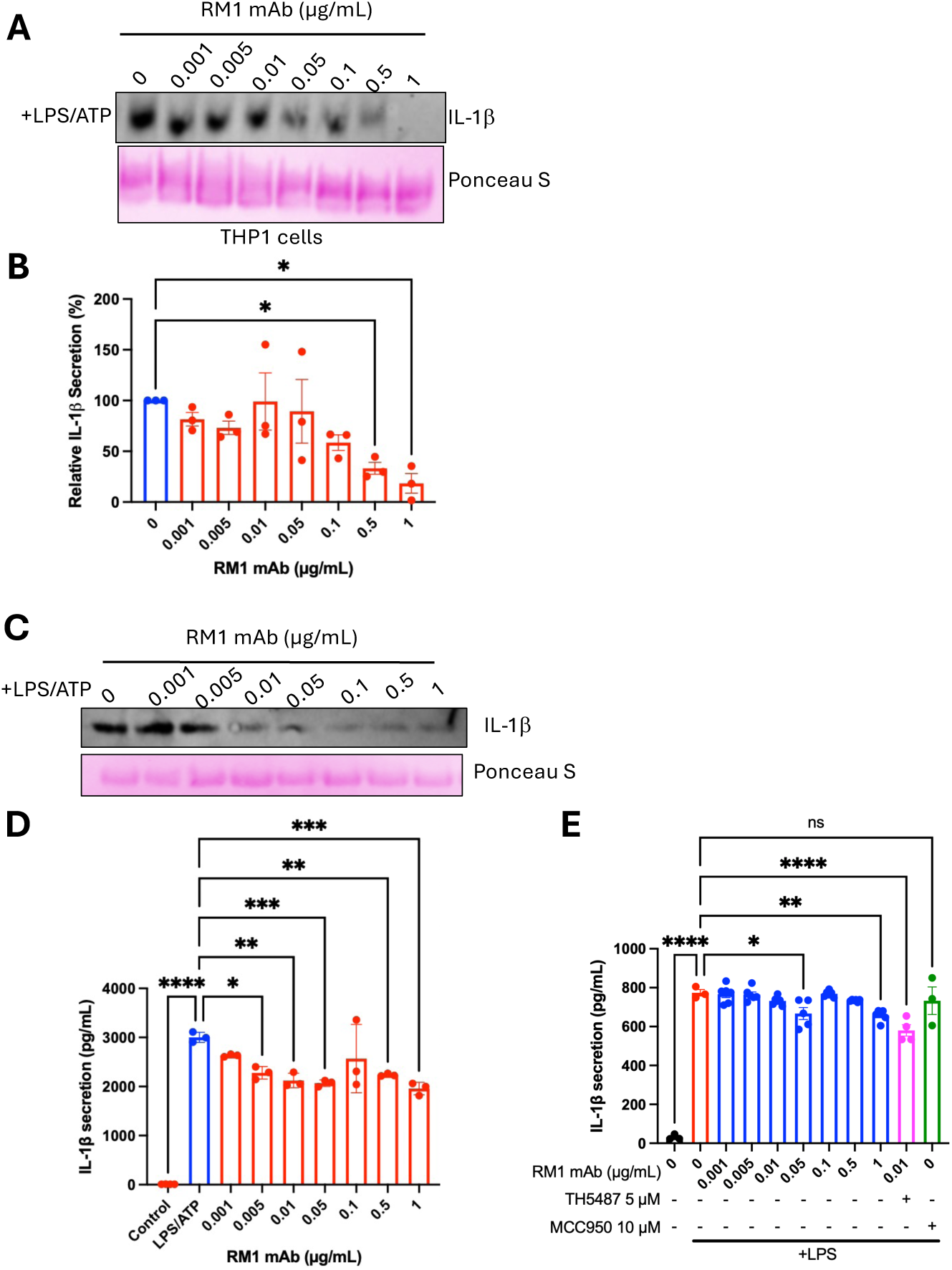
RM1mAb inhibits inflammasome activity in immortalized and primary human cells. A) Immortalized human THP1 cells were primed with LPS, challenged with 0.001-1 µg/mL RM1mAb, and activated with ATP. IL-1β secretion was visualized by western blot. B) IL-1β secretion was quantified by evaluating western blot band intensity, showing inhibition with 0.5 ug/mL RM1mAb. C) Primary human PBMCs were primed with LPS, challenged with 0.001-1 µg/mL RM1mAb, and activated with ATP. IL-1β secretion was visualized by western blot. D) IL-1β secretion was quantified by ELISA E) Primary human PBMCs from CAPS donors were primed with LPS, and challenged with 0.001-1 µg/mL RM1mAb, 5 µM TH5487, or 10 µM MCC950. IL-1β secretion was quantified by ELISA. For all graphs, N=3, error bars signify mean +/- SEM. Analyzed by one way ANOVA where *P<0.05, **P<0.01, ***P<0.001, ****P<0.0001

## Discussion

Our results validate that structural biology is a valuable tool to intuitively design monoclonal antibodies to inhibit cytosolic targets in inflammasome activated cellular states. We designed a full human IgG1 containing two Fabs capable of interacting with NLRP3 pyrin domain (**Fig. S4, S5**). *In silico* support for RM1mAb utilizing FcγR1 to inhibit the interaction of NLRP3 with ox-mtDNA and subsequent inhibition of inflammasome activation was validated herein with human monocytes and FCAS PBMCs.

CD64 expression (FcγR1) is IFN-inducible in monocytes of patients with systemic lupus erythematosus(*36*). Inhibition of NLRP3 with TH5487 decreases IL-1β and inflammasome formation and simultaneously increases in pro-survival IFN-β in human PBMCs(*34*). We demonstrate biological targeting of NLRP3 via its pyrin domain with RM1mAb. FcγR-mediated IgG internalization would normally result in antibody destruction via lysosomes (*33*) or recycling back to the cell surface. Herein, we exploit the fact that NLRP3 activators promote lysosomal disruption(*33*) and use FcγRs to internalize IgG’s that inhibit IL-1β secretion (**Fig. 7**). Moreover, low concentrations of RM1mAb and TH5487(*7*), which both target the NLRP3 pyrin domain, synergistically inhibit inflammasome activation at concentrations where either alone is ineffective. We posit enhanced synergy of TH5487-mediated increase in IFN(*34*), along with a corresponding increase in FcγR-1(*36*), enable more efficient internalization of RM1mAb which supports high efficacy. Moreover, these biological and chemical-based inhibitors both prevent NLRP3 from interacting with oxidized mitochondrial DNA, which is a key mediator inflammasome activation and subsequent systemic release.

**Figure 7:**
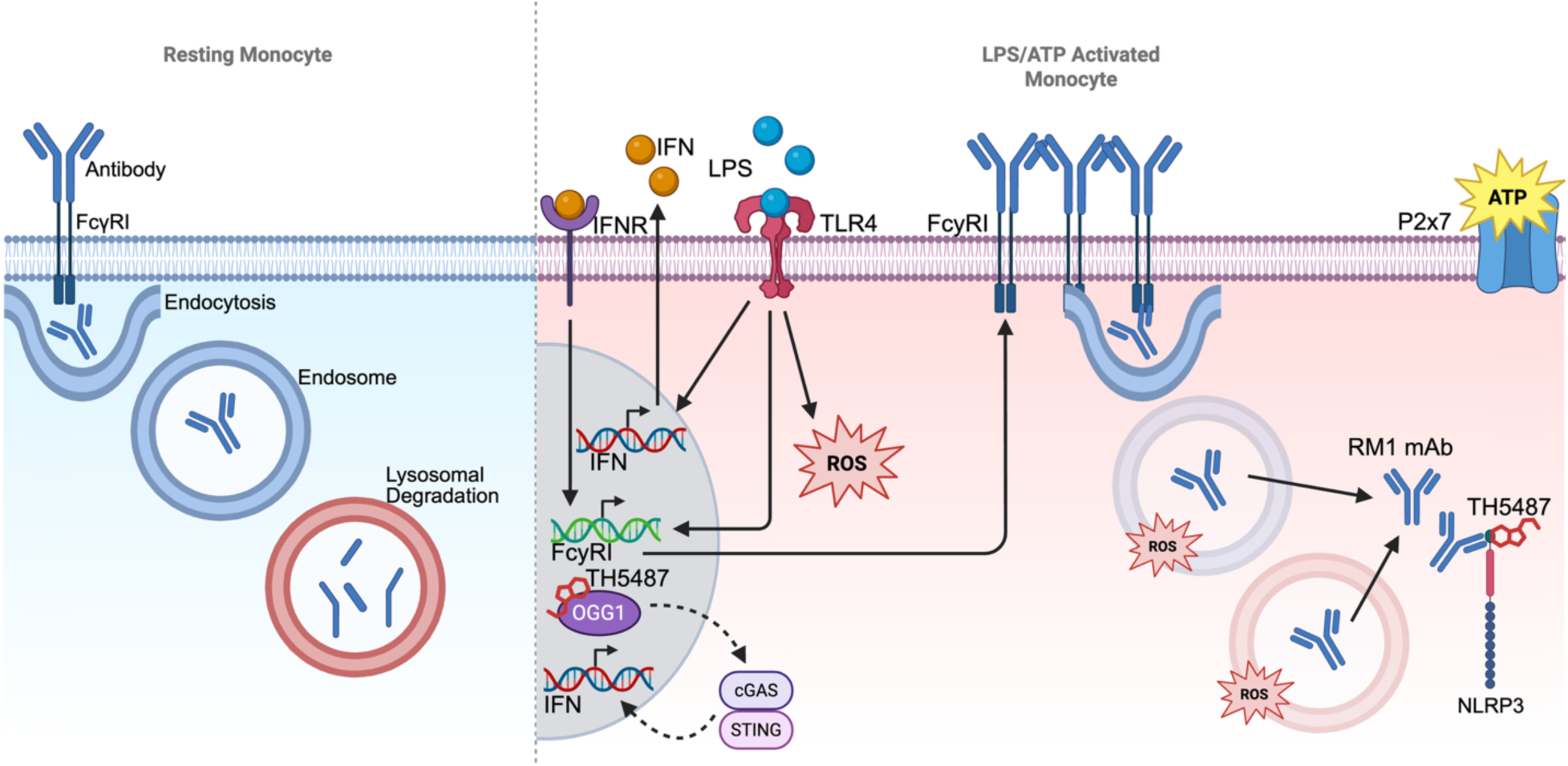
Graphical summary of the mechanism for RM1mAb internalization and synergistic NLRP3 inhibition with TH5487.

LPS-induced endotoxemia-stimulated monocytopoiesis results in rapid expansion and release of new monocytes from bone marrow to circulation in response to systemic LPS exposure in a type 1 IFN-dependent manner(*37*). These new monocytes undergo NLRP3 inflammasome mediated pyroptosis, adding additional mtDNA to circulation and further exacerbating systemic inflammation. Extracellular oxidized mitochondrial DNA itself can bind IgGs and become internalized via FcγR1, which promotes NLRP3 inflammasome activation and systemic inflammation(*38*). In diseases where monocytes provide a major fuel for inflammation, such as Myelodysplastic syndromes (MDS), CAPS, sepsis, and rheumatoid arthritis, inhibiting monocytopoiesis could be useful in preventing cytokine storms. This also extends to several cancers where tumor-associated macrophages play a role involving NLRP3 including pancreatic ductal adenocarcinoma(*39, 40*) and chronic hepatocellular carcinoma(*41*). Future work will elucidate the 3D structure of NLRP3 with oxidized DNA and antibodies/inhibitors described herein(*42, 43*).

## Methods

### De Novo Anti-Pyrin NLRP3 Antibody Design

De novo antibodies targeting the NLRP3 PYD were designed using the RFantibody pipeline, an antibody-specific adaptation of RFdiffusion, as previously described (*18, 44*). Ten structural backbones for variable heavy (VH) and variable light (VL) regions were generated using RFantibody’s antibody-finetuned RFdiffusion component, with NLRP3 PYD hotspot residues specified as binding targets through the protein-protein interaction interface design module. The generated backbones contained poly-glycine placeholder sequences in the CDRs, which were subsequently designed using RFantibody’s ProteinMPNN interface design module configured for antibody sequence optimization. Three sequence variants were generated per structural backbone using a sampling temperature of 0.1 to balance sequence diversity with design quality, yielding 30 total antibody designs with realistic amino acid compositions in the CDR loops while preserving framework region integrity. Structural validation and binding prediction were performed using RF2, the antibody-trained version of RoseTTAFold2 that incorporates antibody-specific structural knowledge, with confidence scores calculated across all atom positions. The top three candidates were selected based on a composite scoring function incorporating RF2 confidence scores, binding interface geometry, sequence diversity metrics, and absence of structural clashes as evaluated through PyRosetta energy calculations. To generate full-length therapeutic antibodies suitable for mammalian expression, the optimized VH and VL variable regions were grafted onto human IgG1 constant domains, also using PyRosetta, to perform sequence-structure alignment with an IgG backbone (PDB: 1IGY), replacing only the variable regions while preserving the constant domain architecture. Final antibody-antigen complex structures were validated using AlphaFold3 to confirm maintained binding interactions and overall structural integrity. For experimental validation, separate expression constructs were designed for heavy and light chains, each incorporating optimized Kozak consensus sequences and secretion signal sequences to enable co-expression, proper folding, and secretion in mammalian cell systems with subsequent disulfide-mediated assembly into functional IgG antibodies. These designs were sent for DNA expression by Twist Biosciences.

### Molecular Dynamics

#### Modeling of the NLRP3-RM1mAb complex

The all-atom model of the NLRP3-antibody complex was constructed from the AlphaFold3 model an NLRP3 monomer PDB 7PZC and bound to RM1mAb. The complex was assembled using Visual Molecular Dynamics (VMD)(*45*) and parameterized using the all-atom additive CHARMM36m force field for protein(*23*). The system was solvated with TIP3P explicit water molecules in a rectangular box with at least 15 Å between the protein and box edges, and the charge was neutralized with Na⁺ and Cl⁻ ions up to a concentration of 150 mM. The final system accounted for approximately 200,000 atoms.

#### All-atom MD simulations of NLRP3-RM1mAb complex

Conventional all-atom MD simulations of the NLRP3-Am1mAb complex were performed using NAMD 2.14 (*22*) as previously described(*46, 47*). Energy minimization was initially performed for 10,000 steps utilizing the conjugate gradient method. During this phase, protein backbone atoms were subjected to positional restraints, applying a force constant of 1.0 (kcal/mol)/Å². Following this, a 0.5 ns NVT equilibration (including the same harmonic restraints) was conducted. Next, restraints were released, and an additional 5 ns of NPT (isothermal-isobaric) equilibration was performed. Subsequently, NPT production MD simulations were run for 50 ns. Control of the temperature and pressure was achieved with a Langevin thermostat (310 K) and a Nosé-Hoover Langevin barostat (1.01325 bar). The Langevin thermostat was applied with a damping coefficient of 1 ps⁻¹, and the Langevin piston was controlled with a period of 200 fs and a decay time of 100 fs. All simulations were performed using an integration time step of 2 fs and the SHAKE algorithm(*48*) to keep covalent bonds involving all hydrogen atoms fixed. Periodic boundary conditions were active, with the particle-mesh Ewald method(*49*) used for long-range electrostatics calculation with a grid spacing of 1.0 Å. Non-bonded interactions, such as van der Waals interactions and short-range electrostatics, were accounted for with a cutoff of 12 Å and switching distance of 10 Å. Trajectory frames were saved every 10 ps for subsequent analysis.

#### Simulation analysis

Structural stability was assessed by calculating root mean square deviation (RMSD) of Cα atoms relative to the initial equilibrated structure. Protein-protein interactions at the binding interface were characterized through contact map analysis performed with PyContact(*50*), monitoring the number of interprotomer residue-residue contacts within 5 Å between NLRP3 and RM1mAb chains throughout the simulation. Hydrogen bonds were identified using distance and angle cutoffs of 3.5 Å and 30°, respectively. Per-residue contact frequency maps were generated to identify persistent interaction hotspots at the binding interface, with contact frequencies calculated relative to the total number of frames analyzed. Root mean square fluctuation (RMSF) analysis was performed on Cα atoms to quantify residue-specific flexibility after alignment to the reference frame. Center of mass distances between NLRP3 and the antibody were monitored to assess complex stability over the simulation trajectory. All trajectory analyses were performed using VMD, PyContact, and Tcl scripts. Molecular visualizations were generated with VMD.

### NLRP3 Purification

NLRP3 purification was performed as previously described(*6, 7*) DNA for wild-type full-length NLRP3 was expressed in DH5α cells and purified using the PureLink HiPure Plasmid Maxiprep Kit (Thermo Fisher). The protein was expressed using the Expi293 Expression System (Thermo Fisher) per the manufacturer’s instructions. Cells at 3×10^6^ cells per milliliter were transfected with 1 μg of expression vector per 1 mL of cells. ExpiFectamine 293 Enhancer 1 and Enhancer 2 were added approximately 16 hours post-transfection Once the cells reached viability of ≤80% live cells (usually 4-5 days), they were pelleted by spinning at 300 rpm for 5 min at 4 °C. The cells were washed with cold PBS and spun again at 1200 rpm for 5 min. The pellet was then lysed with 50 mM Tris pH 7.4, 1 mM PMSF, 1 x protease and phosphatase inhibitor, 300 mM NaCl, 0.1% SDS, 10% glycerol, and 1% Triton X-100 by sonication for 42 s in intervals of 2 s on, 8 s off. The lysate was further clarified by spinning at 100,000 × g for 60 min at 4 °C. The supernatant was passed through a 0.45 μm filter and applied to a HisTrap FF crude 5 mL column equilibrated with wash buffer (20 mM Tris, 200 mM NaCl, 10% glycerol, 1 mM DTT, and 25 mM imidazole, pH 7.4). After the sample was loaded onto the column, it was washed with 10 column volumes of the wash buffer and eluted using 20 mM Tris, 200 mM NaCl, 10% glycerol, 1 mM DTT, and 250 mM Imidazole at pH 7.4. Peak fractions further purified by size exclusion using a HiLoad 16/600 Superose 6 pg size exclusion column equilibrated in 20 mM Tris, 200 mM NaCl, 10% glycerol, and 1 mM DTT, pH 7.4. Peak fractions were concentrated using an Amicon Ultra-15 centrifugal filter units (100 kDa MWCO) and analyzed by SDS page and western blot for purity.

### RM1mAb Expression and Purification

Expi293 cells (Thermo Fisher) were transfected with DNA for the heavy chain and light chain of RM1mAb. In order to express both plasmids, 0.5 µg/mL each of heavy and light chain plasmids (1:1 molar ratio) was co-transfected into Expi293F cells maintained at 3×10^6 cells/mL with >95% viability in Expi293 Expression Medium at 37°C with 8% CO2 and 125 rpm orbital shaking. ExpiFectamine 293 Enhancer 1 and Enhancer 2 were added approximately 16 hours post-transfection. The expression cultures were harvested once viability dropped below 80% (typically 5-7 days post-transfection). Upon harvesting, the cultures were centrifuged 3,000×g for 10 minutes at 4°C to pellet cells, the supernatant was retained, and the cell pellet was discarded. The supernatant was filtered through a 0.22 μm membrane filter to remove cellular debris. A HiTrap MabSelect SuRe 5 mL protein A affinity column (Cytiva) was equilibrated with 5 column volumes of phosphate-buffered saline (PBS, pH 7.4) at 4°C prior to use. The filtered supernatant was loaded onto the column at 2.5 mL/min flow rate using an ÄKTA purification system, monitoring A280 absorbance to track protein binding. Unbound proteins were washed away with 10 CV PBS at 5 mL/min until A280 baseline was achieved, then bound protein were eluted using 100 mM sodium citrate buffer (pH 3.0) at 2.5 mL/min, collected in 1 mL fractions. The acidic eluate was immediately neutralized by adding 100 μL of 1 M Tris (pH 9.0) to each fraction containing protein. Peak fractions were pooled and concentrated using Amicon Ultra-15 centrifugal filter units (100 kDa MWCO) while simultaneously performing a buffer exchange into sterile PBS containing 10% glycerol for long-term storage. Determination of the final protein concentration was performed using an A280 nanodrop and the purity and proper heavy/light chain assembly were assessed by SDS-PAGE and western blot analysis.

### Antibody Electromobility Shift Assay

Purified NLRP3 and RM1mAb proteins were mixed at a 1:1 molar ratio and incubated for 1 hour at room temperature. In order to observe complex association, the mixture was run under native conditions on Novex™ Tris-Glycine Mini Protein Gels (Thermo Fisher). Samples for RM1mAb alone, NLRP3 alone, and the NLRP3:RM1mAb mixture were mixed 1:1 with 2x laemmli sample buffer (BioRad) without SDS or BME. On order to maintain the integrity of the complex, none of the samples were boiled prior to running the gel. The gels were run with Novex™ Tris-Glycine Native Running Buffer (Thermo Fisher) at 225 volts for 50 minutes. Gels were transferred onto PVDF membranes using Novex™ Tris-Glycine Transfer Buffer (Thermo Fisher) and probed for with an anti-human Fc HRP-linked antibody. Chemiluminescence reactions were performed and the membranes were visualized using an iBright 1500 imager to analyze any supershift in the RM1mAb.

### Enzyme-linked Immunosorbent Assay (ELISA)

Costar 96 Well Half-Area Microplates (Corning) were coated with 50 µL of 4 µg/mL RM1mAb overnight at room temperature. The next day, wells were washed with 150 µL wash buffer (0.05% Tween 20 in PBS, pH 7.2-7.4, 0.2 µm filtered) 3 times. Plates were blocked with 150 µL Reagent Diluent (1% BSA in PBS, pH 7.2-7.4, 0.2 µm filtered) for 1 hour at room temperature. The plate was washed again as described 3 times, and 50 µL of 1mg/mL NLRP3 was added to each well for 2 hours at room temperature. A no-protein control was also included. Three washes were performed again, and 50 µL of a NACHT-targeting anti-NLRP3 Rabbit antibody (Cell signaling: D4D8T) (Detection 1) was diluted 1:1000 in reagent diluent and added to each well for 2 hours at room temperature. A no-detection 1 control was also included. Three washes were performed again, and 50 µL of an anti-Rabbit HRP-linked antibody (Cell Signaling: 7074) (Detection 2) was diluted 1:1000 in reagent diluent and added to each well for 2 hours at room temperature. A no-detection 2 control was also included. Three washes were performed again, and 50 µL 1-Step™ TMB ELISA Substrate Solution (Thermo Fisher) was added to each well in the dark for 20 minutes at room temperature. Next, 25 µL of stop solution was added to each well and mixed thoroughly. The optical density of each well was determined using a microplate reader set to 450 nm, and wavelength correction set to 540 nm. Corrected A450 values were plotted using GraphPad Prism and analyzed by a one-way ANOVA.

### NLRP3-antibody-mitochondrial DNA binding assay

NLRP3 DNA binding assay was performed as previously described(*7*). Dynabeads M-280 Streptavidin (Thermo Fisher) were washed three times with binding buffer (50 mM Tris, 100 mM NaCl, 2 mM MgCl2, 12% glycerol, and pH 7.4) using a DynaMag-2 magnet. Biotinylated 20-basepair ox-mtDNA and non-ox-mtDNA sourced from IDT were diluted 1:100 from the 100 μM stock solution and incubated with the beads overnight at 4°C while rotating. The following day, inhibitors RM1mAb were serial diluted with binding buffer such that the addition of inhibitor at various concentrations was always 50% of the final volume and final concentration from 0.02-200 µg/mL. The antibody was incubated with the NLRP3 rotating at 4°C for 1 h. A Control of protein with binding buffer was also included. During these incubations, the beads with DNA were washed three times with binding buffer to remove any unbound DNA. Once the NLRP3:RM1mAb incubations were complete, each incubation was added to a well with beads in triplicate and incubated at 4 °C for 1 hour. A control of 200 µg/mL RM1mAb with both ox-mtDNA and non-ox-mtDNA bound beads was also included. After the incubation, the beads were washed again three times with binding buffer and resuspended in LDS sample loading buffer and reducing agent (Invitrogen), each at a final concentration of 1X. The samples were boiled at 90°C and run on NuPAGE 4 to 12%, Bis-Tris 1 mM 15-well mini-gels at 200 V for 30 min. Samples were transferred to PVDF membranes probed with an anti-NLRP3 antibody (AdipoGen) at a 1:10,000 dilution in 2.5% BSA in TBST. Blots were incubated with an HRP-linked anti-mouse antibody (Cell Signaling) for 1 h at a 1:1,000 dilution and imaged using the iBright 1500 Imaging system. The intensities of the bands were quantified using the iBright Analysis Software, plotted using GraphPad Prism, and analyzed by a one-way ANOVA.

### Antibody Electroporation Inflammasome Inhibition Assay

THP-1 cells were plated at 1×10^6 cells/mL in RPMI 1640 Medium without phenol red (Thermo) supplemented with 10% heat-inactivated fetal bovine serum (Thermo), 1X MEM Non-essential amino acid solution (Thermo), 1mM Sodium Pyruvate (Thermo), and 1% Penicillin-Streptomycin-Glutamine (Thermo), and allowed to rest for 6-8 hours at 37 °C with 5% CO2. Cells were then primed with lipopolysaccharide (LPS) at a final concentration of 500 ng/mL for 12 hours to upregulate NLRP3 expression and prime the inflammasome pathway. Following LPS priming, cells were washed twice with sterile phosphate-buffered saline (PBS) and resuspended in 9 μL Neon NxT buffer R per well in a 96-well electroporation plate. Anti-NLRP3 antibody or non-targeting Rabbit IgG (Cell Signaling: 2729) solutions were added to achieve the desired final concentrations in a standardized 1 μL volume per well, with antibody concentrations ranging from 1-1000 μg/mL. The 8-channel Neon NxT pipette station (Thermo) was connected to the electroporation device and primed by filling 8-channel tubes with 2 mL of E10 electrolytic buffer, ensuring complete electrode submersion. Sterile 10 μL Neon NxT tips were attached to the 8-channel pipette with careful attention to maintaining electrode contact with the electrolytic buffer. Cell-antibody mixtures (10 μL total volume) were aspirated by pressing the plunger to the first stop, immersing all tips into the sample wells, and slowly releasing to avoid air bubble incorporation. The loaded pipette was docked vertically into the pipette station and electroporation was performed using the following parameters: 1400 V, 20 ms pulse width, and 2 pulses per sample. Immediately post-electroporation, cells were transferred to a pre-warmed 96-well recovery plate containing 100 μL per well of antibiotic-free growth media and incubated at 37 °C with 5% CO₂ for 4 hours to allow membrane recovery and antibody internalization. Following recovery, NLRP3 inflammasome activation was induced by adding ATP to a final concentration of 4 mM for 1 hour at 37°C. Cells were harvested by centrifugation at 500×g for 5 minutes at 4°C, with supernatants collected and stored at -80°C for subsequent IL-1β quantification by ELISA. Cell pellets were washed twice with ice-cold PBS and stored at -80 °C for downstream protein expression analysis by Western blot. The protein concentration of each fraction (supernatant and lysate) was evaluated using a Bradford Assay (BioRad), and western blots and ELISAs were run to quantify protein expression.

### Non-electroporated mediated antibody inhibition assay

THP1 cells were treated as previously described (PMID: 40661406). Briefly, cells at viability >95% and 1*10^6^ cells/mL were split into 24-well TC-treated plates (Corning) with 1 mL of cells per well and 500x lipopolysaccharide (Thermo Fisher) was added to each well for 16 hours for a final concentration of 500 ng/mL. The next day, antibodies were added to each well for 1 hour at 0.001-1 µg/mL. A no-antibody control was also included. Next, LPS-only wells were harvested, and 4 mM ATP was added to each well for 1 hour. The supernatant and cells were separated by spinning at 300 x g for 5 minutes at 4°C. The supernatant fraction was removed from the cell pellet and the cell pellet was resuspended in 1 mL ice-cold PBS, re-pelleted by spinning at 1000 x g for 5 minutes at 4°C, and lysed with 300 µL of RIPA buffer (Boston BioProducts) supplemented with an EDTA-free protease/phosphatase inhibitor cocktail (Roche) by rotating for 5 minutes at 4°C. The lysed cells were spun at 14,000 x g for 15 minutes, and 200 µL of the clarified lysate was removed and saved for analysis. The protein concentration of each fraction (supernatant and lysate) was evaluated using a Bradford Assay (BioRad), and western blots and ELISAs were run to quantify protein expression.

### Primary human PBMC inflammasome activation in the presence RM1mAb

PBMCs from healthy individuals were treated as previously described (PMID: 40661406). Briefly, primary human PBMCs (OrganaBio) were seeded at 2 × 10^6^/mL in 96-well plates in Opti-MEM Reduced-Serum Media (Gibco) and immediately primed with 1.6 µg/mL LPS (Thermo Fisher) for 3 hours. During the third hour, cells were treated with 0.001-1 µg/mL Rm1mAb. A no-antibody control was also included. The LPS-only samples were harvested, and 4 mM ATP was added to remaining samples for 1 hour. Cells were harvested by spinning at 1000 x g for 5 minutes. The supernatant was removed and saved for IL-1β western blots and ELISAs, and the cells were washed 3 times with sterile pre-chilled PBS, lysed with 50 µL of RIPA buffer (Boston BioProducts) supplemented with an EDTA-free protease/phosphatase inhibitor cocktail (Roche) by rotating for 5 minutes at 4°C. The lysed cells were spun at 14,000 x g for 15 minutes and 40 µL of the clarified lysate was removed and saved for analysis. The protein concentration of each fraction (supernatant and lysate) was evaluated using a Bradford Assay (BioRad) for western blot and ELISA analysis.

### FCAS PBMC inflammasome activation in the presence of RM1mAb, TH5487, and MCC950

PBMCs from patients harboring the L353P FCAS mutation were a generous gift of Dr. Hal Hoffman (UCSD). Cells were treated as previously described (PMID: 40661406). Briefly, cells were seeded at 2 × 10^6^/ml in 96-well plates in Opti-MEM Reduced-Serum Media (Gibco). Cells were primed with 1.6 µg/ml LPS (Thermo Fisher) for 3 hours. During the third hour, cells were treated with 0.001-1 µg/mL Rm1mAb, 0.1-100 µM MCC950 (Sigma), or a combination of 0.01 µg/mL RM1mAb and 5 µM TH5487 for 1 hour. A no-antibody/drug control was also included. The cells were harvested by spinning the plates at 1000 x g for 5 minutes. The supernatant was removed and saved for IL-1β western blots and ELISAs, and the cells were washed 3 times with sterile pre-chilled PBS and lysed as described above for concentration normalization, and western blot and ELISA analysis.

### Cell Death Quantification by LDH Measurement

Cell viability following antibody electroporation and NLRP3 inflammasome activation was assessed by measuring lactate dehydrogenase (LDH) release into culture supernatants using the CytoTox 96 Non-Radioactive Cytotoxicity Assay (Promega) according to manufacturer specifications. Cell culture supernatants (50 μL) collected from experimental wells and cell-free medium controls were transferred to a clear 96-well microplate and incubated with an equal volume (50 μL) of CytoTox 96 reagent for 30 minutes at room temperature in the dark to allow enzymatic conversion of tetrazolium salt to formazan product. The colorimetric reaction was terminated by addition of 50 μL stop solution to each well, and absorbance was measured at 490 nm using a microplate reader (SpectraMax, Molecular Devices). Percentage cytotoxicity was calculated using the formula: [(Experimental LDH release - Spontaneous LDH release) / (Maximum LDH release - Spontaneous LDH release)] × 100, where spontaneous release represents LDH from untreated control cells, experimental release represents LDH from antibody-treated cells, and maximum release represents LDH from cells lysed with the provided lysis solution.

### Quantification and statistical analysis

A one-way ANOVA was used to conduct all statistical analyses herein. All statistical analyses were performed as indicated in the figure legends, where N represents the number of replicates. Bar graph data were presented as mean ± SEM. P-values <0.05 were considered statistically significant.

### Data and materials availability

All data are available in the main text or the supplementary materials.

### Study Approval

Studies performed using PBMCs obtained from patients with CAPS received the approval of the University of California Human Research Protection Program committee, and informed consent was obtained from the subjects before the study. Institutional Review Board approval number is 180064.

## Author contributions

Conceptualization: AL, RM

Methodology: AL, VH, KW, SP

Investigation: AL, SP, RM

Funding acquisition: RM

Supervision: RM

Writing – original draft: AL, RM

Writing – review & editing: AL, RM

---

**Figure S1:**
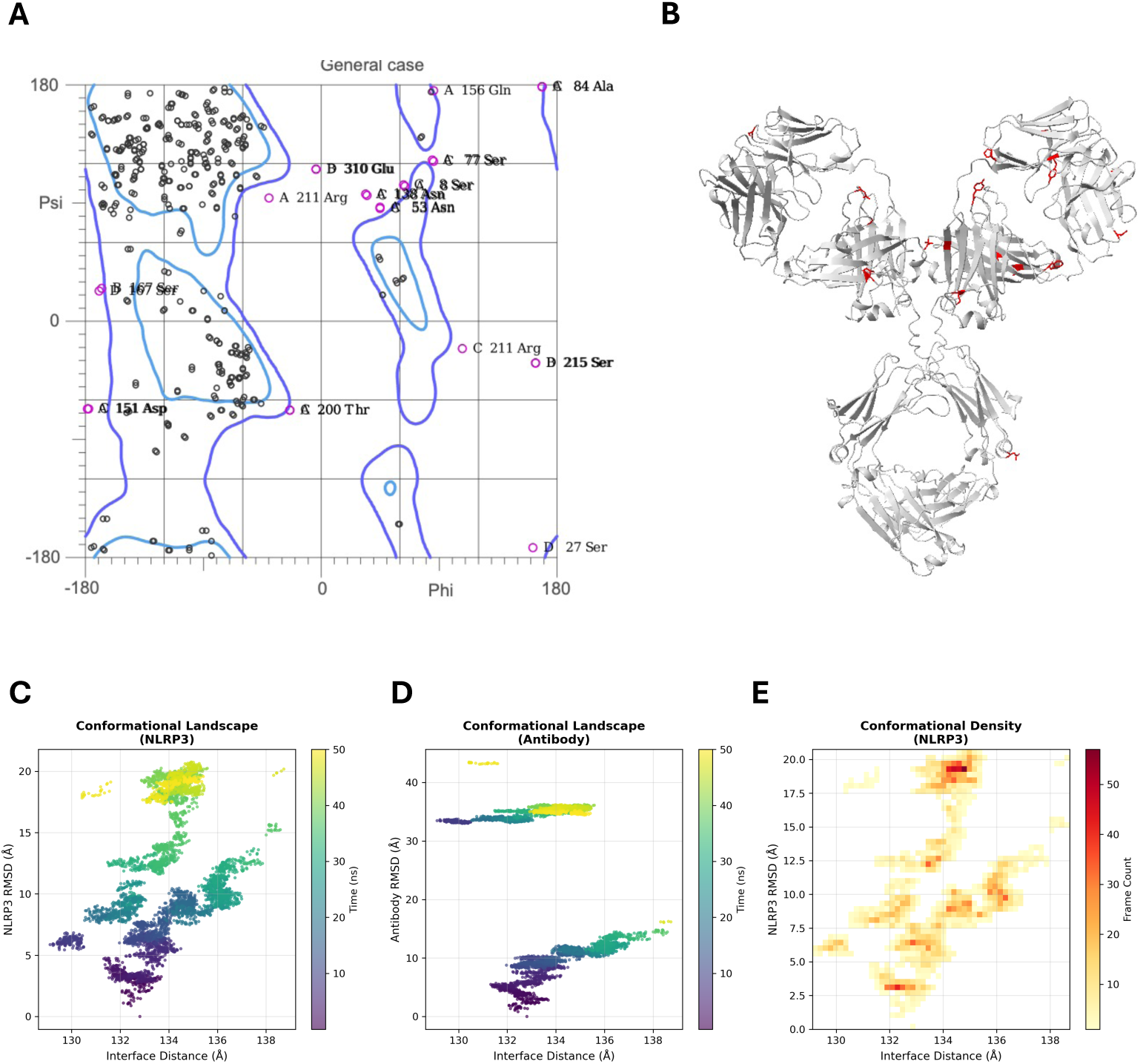
RM1mAb folds into an energetically favorable conformation and binds to NLRP3 at an adaptive interface. A) Ramachandran plot analysis of the structure of RM1mAb showing 78.9% (382/484) of all residues were in favored (98%) regions, and 94.2% (456/484) of all residues were in allowed (>99.8%) regions. B) The structure of RM1mAb (grey) with the residues in disallowed regions highlighted in red, showing their location in loop regions. C) The conformational landscape of NLRP3 over time during a 50 ns molecular dynamics simulation with RM1mAb. Y-axis shows NLRP3 structural changes by RMSD. X-axis shows the interface distances between NLRP3 and RM1mAb in angstroms. Lighter colors indicate later timepoints. D) The conformational landscape of RM1mAb over time during a 50 ns molecular dynamics simulation with NLRP3. Y-axis shows RM1mAb structural changes by RMSD. X-axis shows the interface distances between NLRP3 and RM1mAb in angstroms. Lighter colors indicate later timepoints. E) The conformation density plot of the interface between RM1mAb and NLRP3 during a 50s molecular dynamics simulation. Y-axis shows NLRP3 structural changes by RMSD. X-axis shows the interface distances between NLRP3 and RM1mAb in angstroms. Darker colors indicate maintained positions of the binding interface.

**Figure S2:**
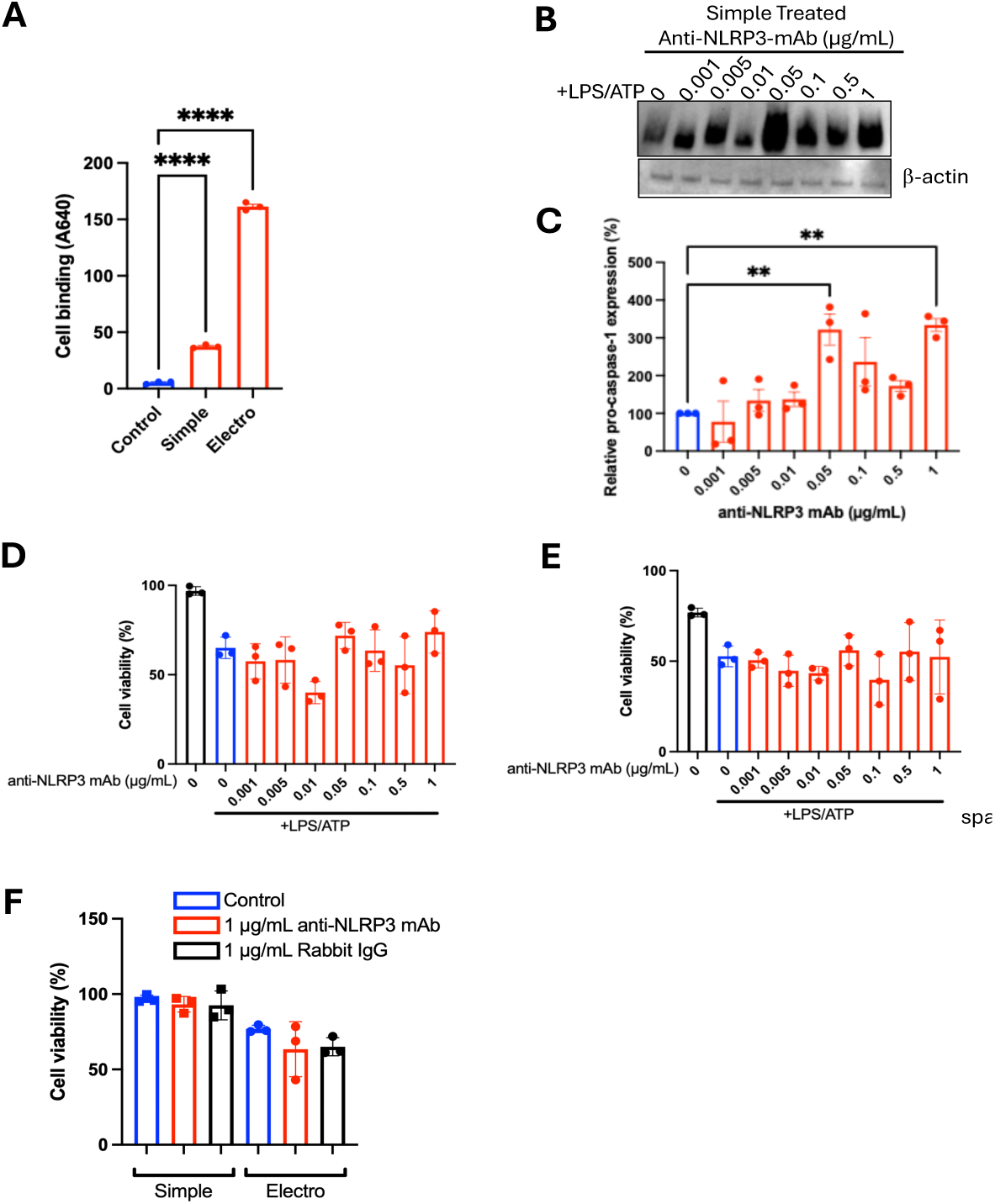
Anti-NLRP3 mAb increased retained intercellular pro-capsase-1 and can enter the cell through simple treatment or electroporation without effecting cell viability compared to LPS/ATP controls. A) Anti-NLRP3 mAb was labeled with AlexaFlour A640 NHS ester chemistry at a 1:1 ratio. The antibody was introduced to immortalized human THP1s by either simple exposure treatment or electroporation. The cells were washed with PBS and the amount of A640 signal on/in the cells was evaluated with a plate reader. B) Immortalized human THP1 cells were primed with LPS, challenged with 0.001-1 µg/mL Anti-NLRP3 mAb, and activated with ATP. Cells were lysed and pro-caspase-1 expression was evaluated by western blot. b-actin was used as a loading control. C) pro-caspase-1 expression was quantified by evaluating western blot band intensity D) Immortalized human THP1 cells were primed with LPS, challenged with 0.001-1 µg/mL Anti-NLRP3 mAb, and activated with ATP. Cell viability was analyzed by LDH release. E) Immortalized human THP1 cells were primed with LPS, challenged with 0.001-1 µg/mL Anti-NLRP3 mAb, and activated with ATP. Control cells were untreated and electroporated with PBS. Cell viability was analyzed by LDH release. F) Immortalized human THP1 cells were treated with anti-NLRP3 mAb by either simple exposure treatment or electroporation. The cell viability was analyzed by LDH release. For all graphs, N=3, error bars signify mean +/- SEM. Analyzed by one way ANOVA where **P<0.01 and ****P<0.0001

**Figure S3:**
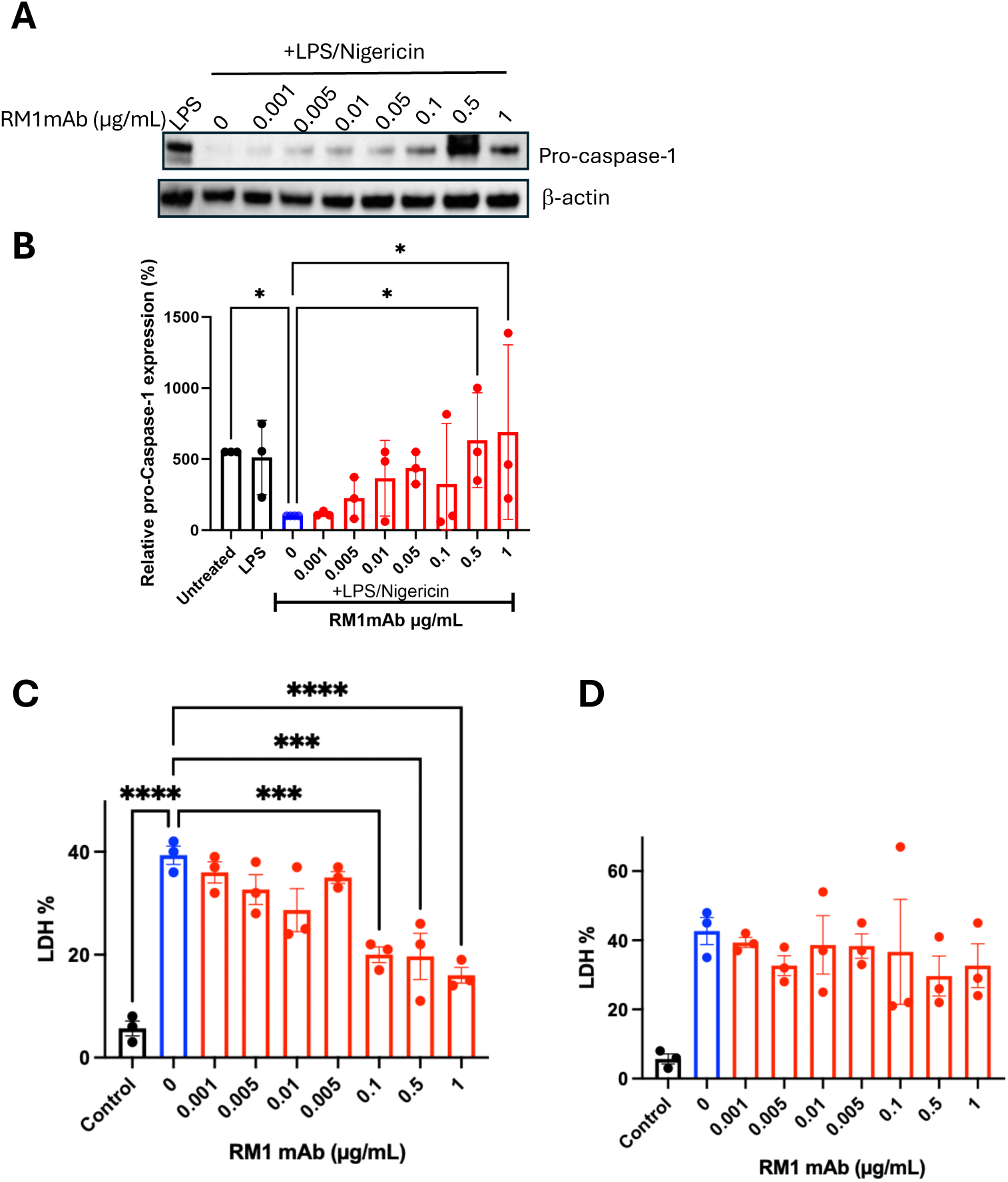
Treatment with RM1mAb increases cytosolic pro-caspase-1 in THP1 cells and reduces cell toxicity in wild-type PBMCs compared to LPS/ATP controls. A) Immortalized human THP1 cells were primed with LPS, challenged with 0.001-1 µg/mL RM1mAb, and activated with Nigericin. Cells were lysed and pro-caspase-1 expression was evaluated by western blot. β-actin was used as a loading control. B) pro-caspase-1 expression was quantified by evaluating western blot band intensity. C) Primary human PBMCs from healthy donors were primed with LPS, challenged with 0.001-1 µg/mL Anti-NLRP3 mAb, and activated with ATP. Cell viability was analyzed by LDH release. D) Primary human PBMCs FCAS patient donors were primed with LPS, and challenged with 0.001-1 µg/mL Anti-NLRP3 mAb. Cell viability was analyzed by LDH release. For all graphs, N=3, error bars signify mean+/- SEM, analyzed by one-way ANOVA with *p<0.05, ***p<0.001, and ****p<0.0001

**Figure S4:**
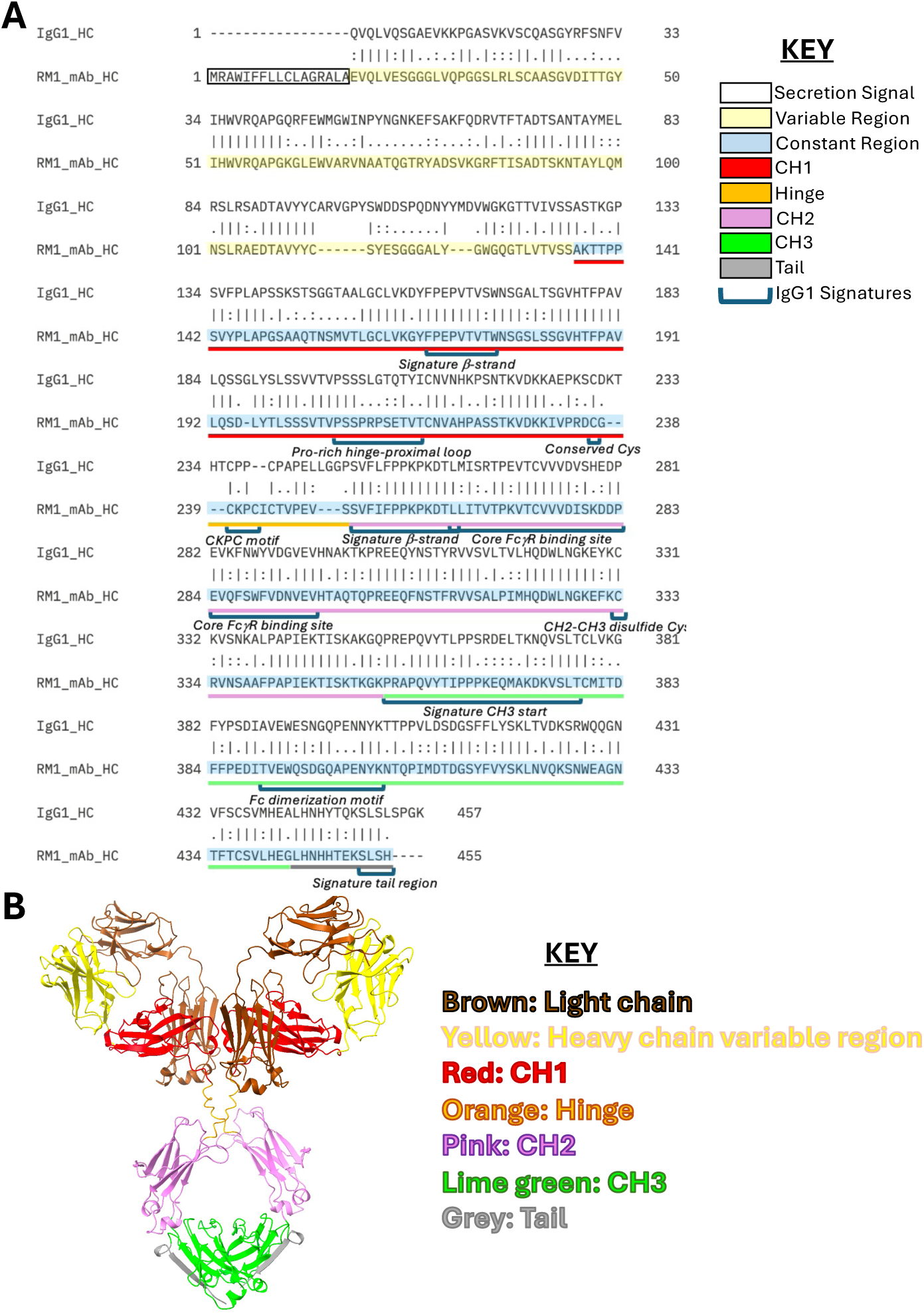
RM1mAb matches the conserved constant region of a human IgG1: A) Sequence alignment of human IgG1 HC (top) and RM1mAb (bottom). Color highlights indicate the secretion signal (white), variable region (yellow) and constant region (blue). Color bars indicate regions in the constant region corresponding to the CH1 (red), hinge (orange), CH2 (purple), CH3 (green) and tail (grey). Noted brackets are conserved regions matching a canonical human IgG. B) Matching structural representation of RM1mAb from (A).

**Figure S5:**
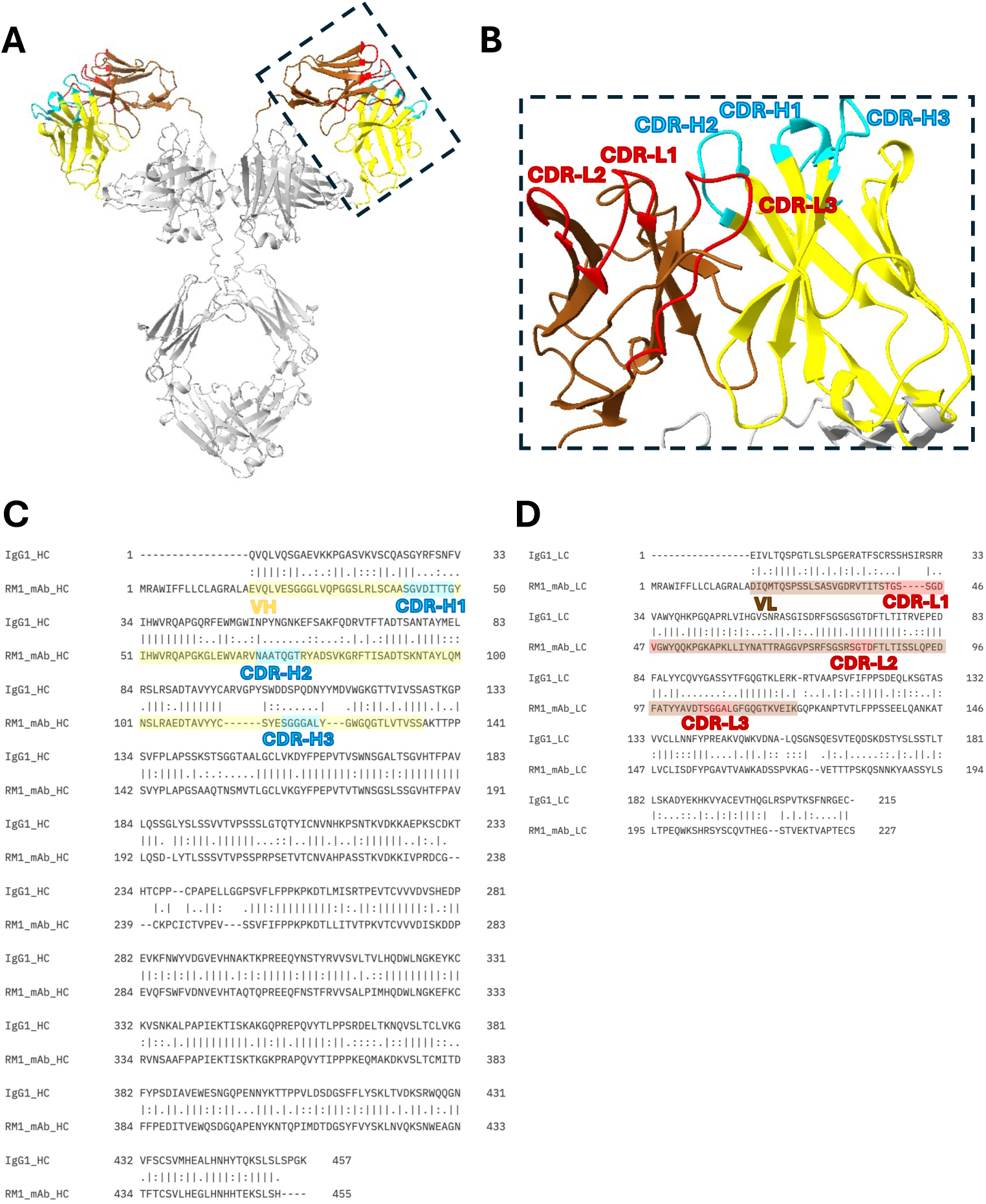
Identification of RM1mAb CDR regions. A) Predicted structure of RM1mAb showing the variable heavy chain (yellow) and heavy chain CDRs (cyan), variable light chain (brown) and light chain CDRs (red), and the constant regions shown in grey. B) Zoomed-in view from dashed box in (A), illustrating variable heavy (yellow residues 1-119) and variable light (brown residues 1-107), CDR-L1 (residues 24-30), CDR-L2 (residues 64-67), CDR-L3 (residues 88-93), CDR-H1 (residues 25-32), CDR-H2 (residues 52-58), and CDR-H3 (residues 100-105). C) Sequence alignment of human IgG1 HC (top) and RM1mAb HC (bottom). Color highlights indicate the variable region (yellow), and CDRs (cyan). D) Sequence alignment of human IgG1 LC (top) and RM1mAb LC (bottom). Color highlights indicate the variable region (brown), and CDRs (red).

**Table S1:** Key residues at the interface between NLRP3 and RM1mAb. 27 NLRP3 residues and 35 RM1mAb residues make significant contact as identified during a 50 ns NAMD molecular dynamics simulation

| **NLRP3 Residue** | **RM1mAb Residue** |
| --- | --- |
| Thr6 | Val2 |
| Pro33 | Leu4 |
| Gln35 | Gly26 |
| Lys36 | Val27 |
| Gly37 | Asp28 |
| Glu64 | Ile29 |
| Glu65 | Thr31 |
| Lys66 | Gly32 |
| Trp68 | Tyr33 |
| Ala69 | Trp47 |
| Val72 | Val48 |
| Trp73 | Arg50 |
| Ile74 | Asn52 |
| Ala76 | Thr55 |
| Ala77 | Gln56 |
| Asn79 | Gly57 |
| Arg81 | Thr58 |
| Try84 | Arg59 |
| Glu85 | Tyr60 |
| Ala87 | Asp62 |
| Lys88 | Ser63 |
| Arg89 | Lys65 |
| Asp90 | Phe68 |
| Glu91 | Arg87 |
| Pro92 | Ala88 |
| Lys93 | Glu89 |
| Trp94 | Asp90 |
|  | Thr91 |
|  | Ala92 |
|  | Val93 |
|  | Tyr94 |
|  | Glu99 |
|  | Gly101 |
|  | Gly102 |
|  | Ala104 |

